# Detecting Misannotated Long Non-coding RNAs with Training Dynamics of Deep Sequence Classification

**DOI:** 10.1101/2020.11.07.372771

**Authors:** Afshan Nabi, Ogun Adebali, Oznur Tastan

## Abstract

Long non-coding RNAs (lncRNAs) are the largest class of non-coding RNAs (ncRNAs). However, recent experimental evidence has shown that some lncRNAs contain small open reading frames (sORFs) that are translated into functional micropeptides. Current methods to detect misannotated lncRNAs rely on ribosome-profiling (ribo-seq) experiments, which are expensive and cell-type dependent. In addition, while very accurate machine learning models have been trained to distinguish between coding and non-coding sequences, little attention has been paid to the increasing evidence about the incorrect ground-truth labels of some lncRNAs in the underlying training datasets. We present a framework that leverages deep learning models’ training dynamics to determine whether a given lncRNA transcript is misannotated. Our models achieve AUC scores > 91% and AUPR > 93% in classifying non-coding vs. coding sequences while allowing us to identify possible misannotated lncRNAs present in the dataset. Our results overlap significantly with a set of experimentally validated misannotated lncRNAs as well as with coding sORFs within lncRNAs found by a ribo-seq dataset. The general framework applied here offers promising potential for use in curating datasets used for training coding potential predictors and assisting experimental efforts in characterizing the hidden proteome encoded by misannotated lncRNAs. Source code is available at https://github.com/nabiafshan/DetectingMisannotatedLncRNAs.

## 1 Introduction

Genome-wide transcriptome analyses have revealed that the vast majority of the human genome is transcribed; but only 2% of the human genome is annotated as protein coding [11]. A considerable fraction of transcripts are annotated as ncRNAs and lncRNAS constitute the largest category of ncRNAs [10]. While lncRNAs studied are known to play vital roles in cellular processes such as regulation of translation, transcription, chromatin modification and mRNA stability [35,3,39], functions of most lncRNAs remain unknown. Moreover, although lncRNAs-by definition-do not code for proteins, recent studies have shown that short the open reading frames (sORFs) within some lncRNAs are translated into micropeptides of a median length of 23 amino acids [26,1,20,7,13,9]. The translation events of lncRNAs were overlooked previously because the open reading frames (ORFs) present in lncRNAs do not meet the conventional criteria of an ORF: that it encodes at least 100 amino acids in eukaryotes [13]. Despite this, recent studies have shown that micropeptides translated from lncRNAs perform vital functions across species, including bacteria, flies and humans [30,28,13]. Therefore, identifying misannotated lncRNAs is a necessary step towards the functional characterization of this large class of transcripts.

Experimental identification of coding transcripts is performed using ribosome profiling (ribo-seq), which involves capturing and sequencing RNA fragments protected by ribosomes [17]. Use of ribo-seq data has revealed many unexpected protein products [5], including sORFs within lncRNAs [18]. However, since ribo-seq data is known to contain false positives [18,19], several computational methods have been proposed to identify true translated ribo-seq fragments. These include FLOSS [17], ORFscore [4] and PhyloP [32,29]. FLOSS [17] relies on the typical length of ribo-seq fragments to determine truly coding ribo-seq fragments. ORFscore [4] relies on the property that translating ribosomes shift by three nucleotides (ribosome phasing),which leads to a characteristic pattern wherein true positive fragments have higher sequencing reads every third nucleotide. PhyloP is used to find true translated ribo-seq fragments by probing conservation across species [32,29]. These computational methods applied over ribo-seq data can be used to find sORFs that are both translated and located within lncRNAs. However, one major limitation of relying on ribo-seq data to identify misannotated lncRNAs is that not all transcripts are likely to be transcribed and translated at a given time point in a given cell. To obtain a complete picture of the misannotated lncRNAs in the genome, different cell types, at different developmental stages, under different environmental conditions need to be sequenced and analyzed. In contrast, the nucleotide sequence of a lncRNA transcript is unlikely to change across cell types and conditions. Therefore, methods to detect misannotated lncRNAs from nucleotide sequences will be useful in assisting experimental efforts and available ribo-seq based computational methods.

Once sufficient coding sORFs have been detected, a dataset containing positive (coding) and negative (non-coding) examples can be built to train models to predict the coding potential of a given sORF. These methods can then be used to assess the coding potential of a transcript. For instance, logistic regression [42] and SVM [37] based models have been proposed to predict the coding potential of a given sORF with sequence length ≤ 303 nucleotides. To determine whether a lncRNA is misannotated by using these methods requires first to extract all possible sORFs in the lncRNA and then assess the coding potential of each of these sORFs. However, while it is possible to predict the coding potential of lncRNA sORFs with these tools, it is impossible to evaluate the performance since the data on which lncRNA sORFs are truly coding is very sparse [42].

Several classical machine learning [23,21,38,41] and deep learning [14,2,6] based models, which focus on longer length nucleotide sequences as input, have also been developed to predict the coding potential of a given RNA. Most of these methods demonstrate very high prediction performance. However, using these, it is not possible to identify lncRNAs that might be misannotated. This is because these models do not incorporate any strategy to deal with misannotated lncRNAs in the underlying training datasets. To ensure that we are not overfitting the models to learn biologically irrelevant decision boundaries, there is a need to find ways to determine possible misannotated RNAs in the underlying training datasets.

We present a framework that leverages deep learning models’ training dynamics to determine whether a given lncRNA transcript in the dataset is misannotated. In particular, we train convolutional neural network (CNN) [24], long short term memory (LSTM) [15], and Transformer [40] architectures to predict whether a given nucleotide sequence is non-coding or coding and use the training dynamics to identify possible misannotated lncRNAs [36]. Our models learn biologically relevant features to distinguish between coding and non-coding RNAs with average AUC scores >91% and identify many misannotated lncRNAs. By generating unsupervised clusters of coding and non-coding RNAs, we observe that there might be a continuity in the embedded space between coding and misannotated lncRNAs. Finally, our results show a significant overlap with previous methods that use ribo-seq data to identify misannotated lncRNAs as well as with a set of experimentally validated misannotated lncRNAs. This work represents the first instance where deep learning model training dynamics are successfully applied to identify misannotated lncRNAs from nucleotide sequences. This approach can be applied to better curate datasets for training coding potential prediction models and can be applied alongside ribo-seq data to identify misannotated lncRNAs with high confidence.

## 2 Methods

### 2.1 The overall framework

The workflow for determining misannotated lncRNAs is described in Figure 1. The main steps are as follows. We train deep learning based sequence classification models that can distinguish coding and non-coding RNAs. Once we establish that models can achieve good performance on the held-out test data, we retrain a final model on all the data and inspect its training dynamics to find possibly misannotated ncRNAs. By focusing especially on the union of the lncRNAs identified as misannotated by all the models, we arrive at a final list of putative misannotated lncRNAs. We compare this list to experimentally validated coding ncRNAs as well as to a ribo-seq dataset. We use unsupervised clustering to find where possibly misannotated lncRNAs are located within the broader RNA clusters. We also study the features used by the deep learning models to make classification decisions. In the following sections we detail the dataset, the sequence classification models trained and the other analysis we conducted.

**Fig. 1:**
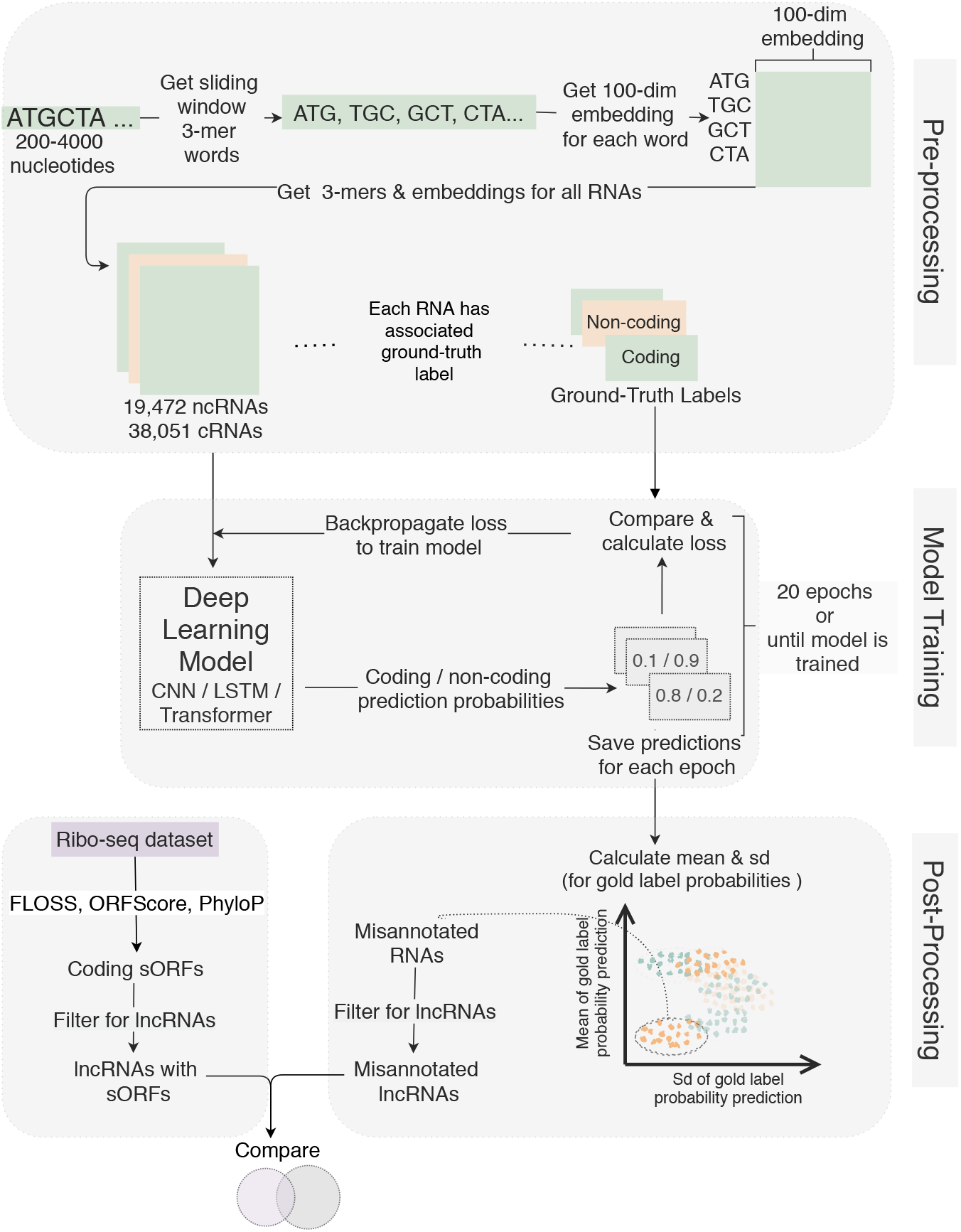
Workflow for identifying misannotated lncRNAs by examining the training dynamics of deep learning models. All RNA sequences are constrained to be between 200-4000 nucleotides long. From each RNA sequence 3-mer ‘words’ are obtained by using a window that slides by 1 nucleotide at each step. For each 3-mer ‘word’, 100-dimensional embeddings [31] are obtained. Each RNA also has an associated ground-truth label, i.e. each RNA is labelled as coding or non-coding. Deep learning models are trained using 100-dimensional embeddings for contiguous 3-mers from the sequences. At the end of each training epoch, the predicted probabilities for each RNA being coding or non-coding are saved. After training, the mean and standard deviation for the ground-truth label probability prediction are calculated and misannotated lncRNAs are identified. These are compared to lncRNAs containing translated sORFs determined from ribo-seq data.

### 2.2 Datasets

We use the dataset of human RNA nucleotide sequences compiled by [38] to train the sequence classification models. After filtering to remove non-coding RNA sequences < 200 nucleotides in length, the data comprises of 38,051 coding RNA and 19,472 non-coding RNA sequences. Filtering non-coding RNAs by length was necessary since the length distributions of coding and non-coding RNAs in the dataset was very different; non-coding RNAs are noticeably shorter than coding RNAs. For the deep learning models to learn biologically relevant features in order to distinguish between coding and non-coding RNAs, equalizing the sequence length distributions was necessary. If the sequence lengths of ncRNAs are significantly shorter than those of coding RNAs, then sequence length itself might be used by the models as a feature distinguishing between coding and non-coding RNAs.

### 2.3 Deep Learning Model Architectures

We train CNN [24], LSTM [15] and Transformer [40] models to classify non-coding and coding RNA sequences. Each input sequence is truncated to a length of 4000 nucleotides before being input to the deep learning models. The sequences are encoded as 1-nucleotide sliding window 3-mers using the 100-dimensional DNA-embeddings generated by [31]. All three models are implemented using Keras [8]. We use ReLu as the activation function. We trained all models to minimize the sparse categorical cross-entropy loss using the Adam optimizer [22]. In all cases, we use a batch size of 64.

#### Convolutional neural network

For the CNN, encoded sequences are fed into an embedding layer which is followed by 3 layers of 1-D convolution (each with 128 units and filter size 5) and max-pooling (5 units). These are followed by a dense layer of 128 units.

#### LSTM

For the LSTM, encoded sequences are fed into an embedding layer which is followed by 2 layers of 1-D convolution (each with 128 units and filter size 5) and max-pooling (5 units), followed by a bi-directional LSTM layer. These are followed by a dense layer of 128 units.

#### Transformer

Encoded sequences are added to a positional encoding and fed into a transformer block followed by global average pooling, dropout and a dense layer of 64 units. The transformer block comprises of a single-headed self-attention layer and a dense layer both followed by layer normalization.

### 2.4 Model Evaluation Set Up

We use the human coding & non-coding train and test datasets provided by [38]. We set aside 20% of the training data as the validation data. We use Keras Tuner [33] to find the optimal set of hyperparameters for the deep learning models. We created a hyperparameter search space for different model architecture and hyperparameter assignment values and used the Hyperband tuner [25] to find the optimal parameters based on validation loss. We tried the following choices for given hyperparameters: dense layer units 64, 128, and 256, 1-D convolutional filters (64 and 128, LSTM units 64, 128, and 256, dropout 0.2, 0.3, 0.4 and 0.5 and learning rate (logarithmic sampling between *e*-2 and *e*-4. We used the best model returned by the Hyperband tuner and retrain a model on the train-validation data to calculate and assess these models’ performances on the held-out test data. Once the test performances are attained, we rebuilt the models on all data to find the misannotated ncRNAs.

Since the training dataset is imbalanced in favor of coding RNA, we used class-weights inversely proportional to the number of class samples to ensure learning. Moreover, since a coding RNA is unlikely to be misannotated, we penalized coding RNA misclassifications 5 times more than non-coding RNA misclassifications.

### 2.5 Identifying misannotated lncRNAs using training dynamics

We inspect the deep learning models’ training dynamics to find possible misannotated lncRNAs. Swayamdipta et al. [36] report that it is possible to identify possibly mislabelled training samples in a given dataset by inspecting how model predictions for samples behave during training. We employ this strategy; at the end of each training epoch, the deep learning models are evaluated on the training examples and the predictions for the class probabilities are saved. Consider a training dataset of size 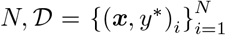 where the *i* th instance consists of the observation, *x_i_* and its true label under the task, *y_i_*\*. We calculate the mean and the standard deviation of the posterior probability of the ground-truth label for example *i* over *E* epochs as follows [36]:

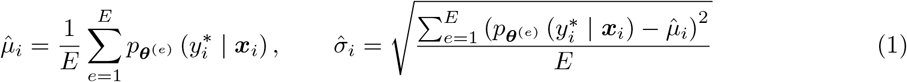

where *p_θ_*(*e*) denotes the probability assigned at the end of the *e*th epoch by the model parameterized with *θ*^(*e*)^. Using the mean and the standard deviation of the predicted probability of ground-truth class across all epochs, the training dataset can be divided into three groups: easy-to-learn, ambiguous, and hard-to-learn. The hard-to-learn samples are those with low mean and low standard deviation of the true class probabilities. In other words, the model consistently misclassifies these samples across training epochs. We retrain the models using both the training & test data and consider the lncRNAs within this hard-to-learn class as candidates for misannotation.

### 2.6 Comparison to cncRNAdb-a manually curated list of experimentally validated coding ncRNAs- and ribo-seq based methods to identify coding sORFs within lncRNAs

We downloaded data from the cncRNAdb [16], a resource that provides a manually curated list of experimentally validated ncRNAs found to be coding. We filtered data to get lncRNAs found to be coding in *Homo sapiens* and compared the list to the misannotated lncRNA candidates generated from the deep learning models.

Next, we compared the list of misannotated lncRNAs generated by our models to a ribo-seq dataset. We downloaded data on sORFs identified in the ribo-seq data generated by [12] from sORFs.org [32]. This database provides computations of values of FLOSS [17], ORFscore [4] and PhyloP [29] metrics for RNAs identified from the ribo-seq data. We used RNAs annotated as lncRNAs and present in both the sequence dataset (used to train deep learning models) and the Ribo-seq dataset in our analysis. According to previous considerations, to get the list of lncRNAs containing translated sORFs, we used the following cutoff values: ‘Good’ for the Floss-classification, ORFscore > 6 and PhyloP > 4 [32].

## 3 Results

### 3.1 Prediction performance of classifying coding vs. non-coding RNAs

Prediction performances calculated on the held-out test set for the models trained are provided in Table 1 and show that our models perform well on the classification task. The LSTM model achieves the highest classification performance with 94% AUC and 96% AUPR. The CNN model performs similarly well with 93% AUC and 95% AUPR, while the transformer achieves 91% AUC and 93% AUPR. Since our aim is to study the underlying dataset and find misannotated lncRNAs, higher prediction performance is not the primary focus. Instead, since we know that the training dataset contains lncRNAs that have incorrect ground-truth labels, we want to ensure that the models are not being overfitted to learn features that might not be relevant to learning the biological distinction between coding and non-coding RNAs as encoded in the nucleotide sequences. In the following sections, we detail how we employ these models to discover possibly misannotated lncRNAs in the underlying dataset.

**Table 1:**
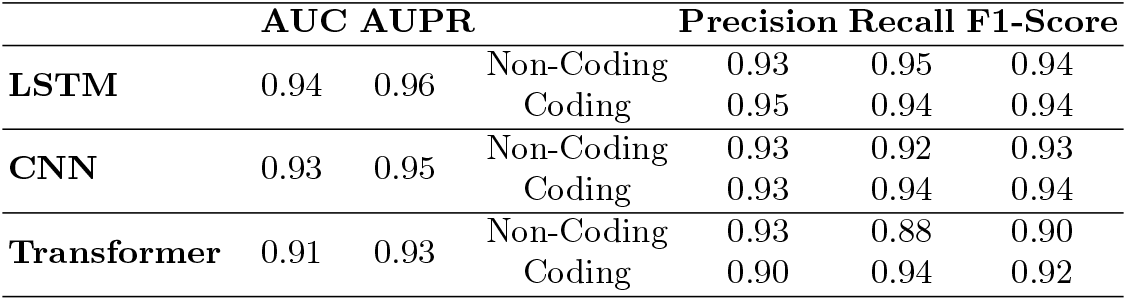
The test-data performances of the different models trained to classify long non-coding RNAs and coding RNAs. AUC and AUPR are micro-averaged.

### 3.2 Training dynamics of deep learning models can be used to identify misannotated lncRNAs

Having evaluated the CNN, LSTM and Transformer models to distinguish between coding RNA and non-coding RNA, we retrain the models using all data and inspect each instances training dynamics. During the training phase of each model, we track the coding probability predictions for each RNA. Figure 2a shows the predictions for the coding probability for three different RNAs across all training epochs for the LSTM model. For example, the coding probability predictions for ENST00000447563 (shown in orange)-an RNA annotated as long non-coding (ground-truth)-are consistently high. In other words, as model training progresses, this RNA is invariably classified as coding. It was recently shown that ENST00000447563 has been misannotated as lncRNA when it can, in fact, code for a protein [13]. Two other examples of correctly annotated coding and non-coding RNA are also shown in Figure 2a. By studying the predictions made by models as they are under training, it is possible to identify putative misannotated lncRNAs.

**Fig. 2:**
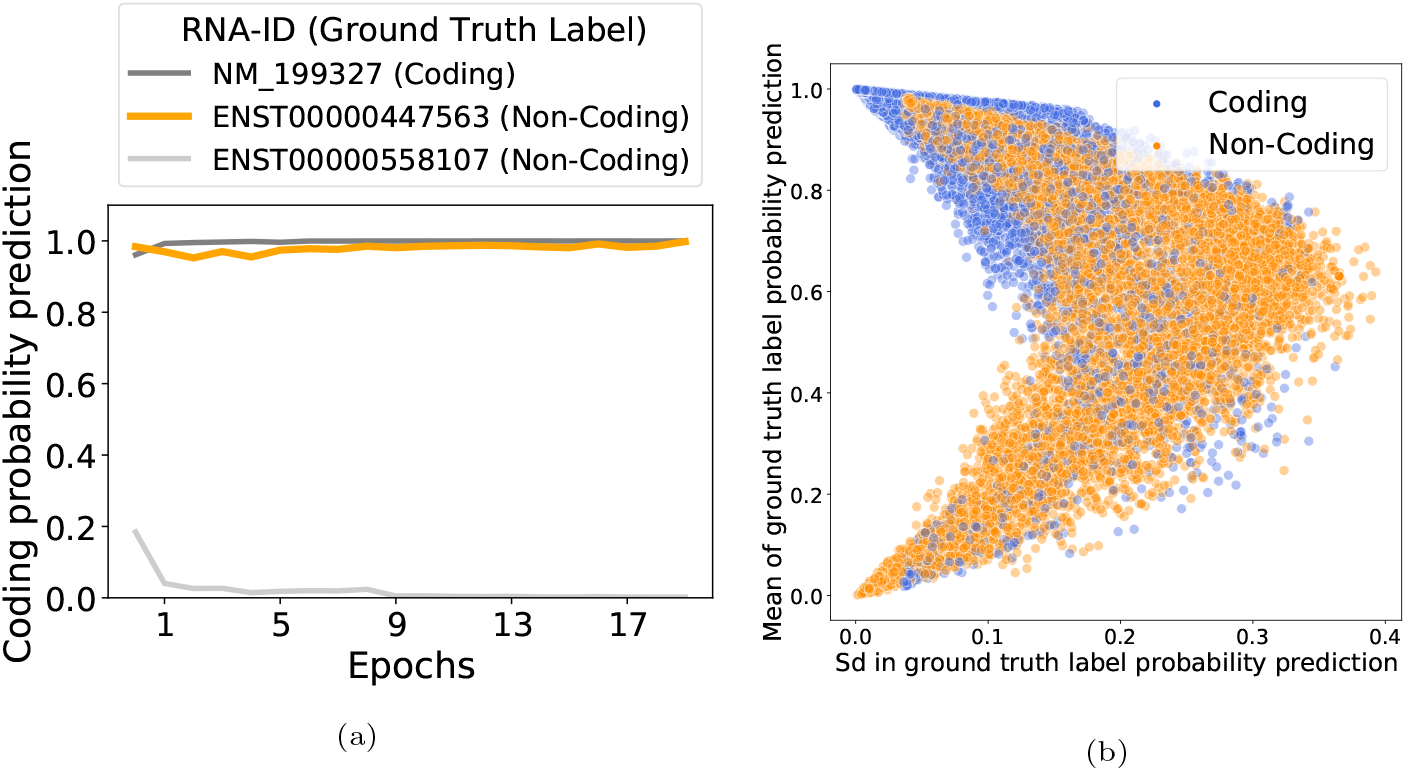
Training dynamics of deep learning models can be used to identify misannotated lncRNAs. (a) Coding probability prediction (probability of being a coding RNA) across all training epochs shown for three RNAs. NM 199327 (ground-truth ‘Coding’) and ENST00000558107 (ground-truth ‘Non-coding’) have high and low coding probability predictions respectively. In contrast, ENST00000447563 consistently has high coding probability prediction, despite having the ground truth-label ‘Non-coding’. This suggests that it might be a misannotated lncRNA. In support of this observation, ENST00000447563 (also known as linc00689) was recently found to be protein coding [13]. (b) Mean (y-axis) and standard deviation (x-axis) of true class label probability predictions across all training epochs can be used to determine misannotated RNAs. The misannotated lncRNAs are those in the bottom left quadrant i.e. lncRNAs with low mean and standard deviation for the ground-truth class (non-coding) probability.

Figure 2b expands upon this idea: calculating the mean and standard deviation of predicted probability for the ground-truth class across all training epochs provides a measure of identifying misannotated lncRNAs. lncRNAs in the lower left quadrant of Figure 2b are considered putative misannotated lncRNAs; these samples have low mean and standard deviation for the predicted probability of the ground-truth class over all training epochs. In other words, these lncRNAs are consistently classified into the non-ground-truth class (coding) and therefore, are likely to be misannotated. It is interesting to note that the majority of the putative mislabelled samples have ground-truth label ncRNA. This observation supports the notion that the current method for identifying putative misannotated lncRNAs is reasonable. This is because an RNA with ground-truth ‘coding’ is unlikely to be misannotated. In conclusion, many lncRNAs might be misannotated and sequence information combined with training dynamics of deep learning based classifiers might help identify such misannotations.

### 3.3 Different deep learning architectures find common misannotated lncRNAs

Figure 3a shows the overlap between the lists of misannotated lncRNAs generated by CNN, LSTM and Transformer models. It is interesting to note that despite the difference in network architectures, the intersection of possible misannotated lncRNAs is large. The CNN model identifies the smallest number of candidate misannotated ncRNAs. It is interesting to note that the number of common candidates identified by Transformer and LSTM but not by CNN (1394 in total) is large as compared to the common candidates between CNN & Transformer only (71) and between LSTM & CNN only (171). 1251 candidates are identified by all 3 models.

**Fig. 3:**
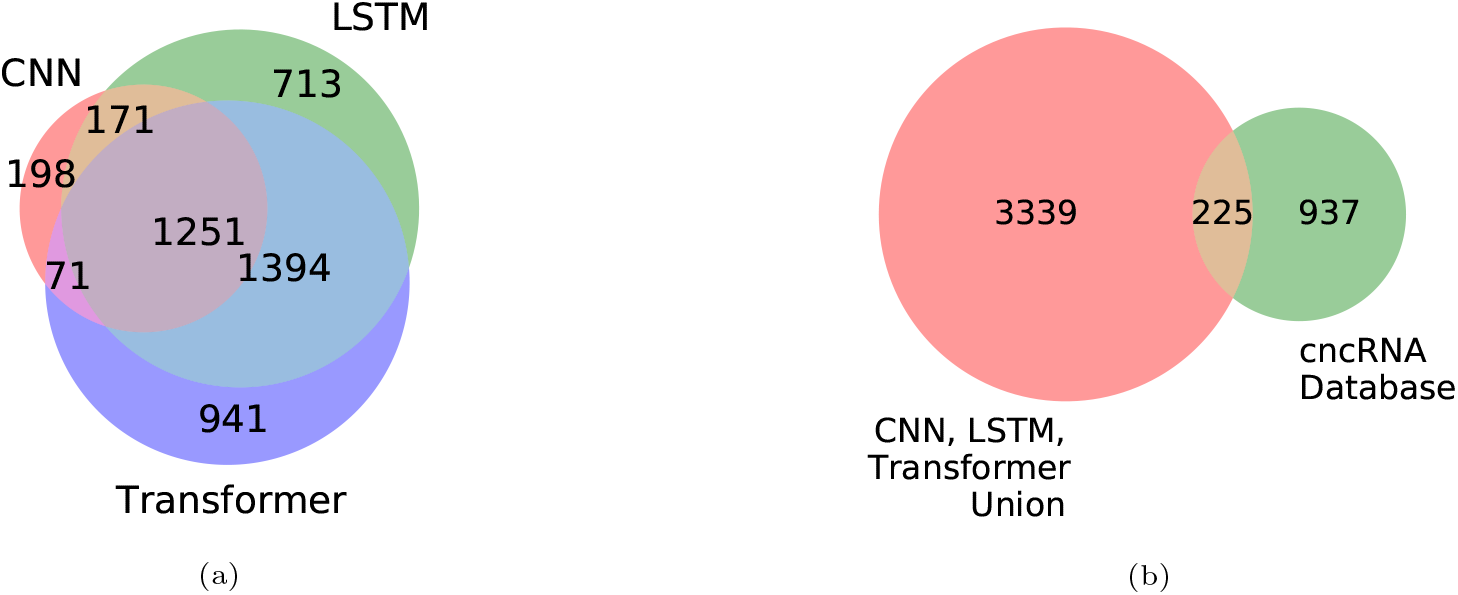
(a) Comparison of the misannotated lncRNAs obtained from by CNN, LSTM and Transformer models’ training dynamics. (b) Comparison of the misannotated lncRNAs obtained from deep learning models with the cncRNA database (hypergeometric test, p-value 1*e*-6), which provides a manually curated list of experimentally validated coding lncRNAs.

### 3.4 Misannotated lncRNAs overlap significantly with manually curated, experimentally validated coding lncRNAs & with misannotated lncRNAs discovered by ribo-seq

To check if the candidate list of misannotated transcripts overlap with already reported misannoated ncRNAs, we find the overlap with the cncRNA database. The cncRNA database provides a manually curated list of experimentally validated coding lncRNAs [16]. Figure 3b shows the overlap between the list of misannotated lncRNAs generated by our deep learning models and the cncRNA database [16]. There are 225 common misannotated lncRNAs, this overlap is highly significant (hyper-geometric test, p-value (1*e*-6)).

Next, we compared the overlap between the misannotated lncRNAs discovered by our deep learning models with a high-throughput ribo-seq dataset. Figure S4 shows the counts for lncRNAs obtained by applying 3 different methods (FLOSS, ORFScore and PhyloP) to identify true positives from ribo-seq data generated by [12]. For FLOSS, lncRNAs with a classification of ‘Good’ are considered candidate misannotated lncRNAs; it is interesting to note that most of the lncRNAs have a ‘Good’ FLOSS score. In contrast, fewer lncRNAs are considered misannotated according to ORFScore and PhyloP. The overlap between these 3 methods to find sORFs from ribo-seq data is shown in Figure S3.

It is important to note that the dataset used in the current work is much smaller and contains fewer lncRNAs than those found from the [12] ribo-seq dataset. In order to be able to compare the numbers of misannotated lncRNAs found by the different methods, we first generated a list of lncRNAs that were present both in the ribo-seq dataset and in the nucleotide sequence dataset used for training deep learning models. From this common lncRNAs master list, we calculated the overlap between misannotated lncRNAs found by different methods. Figure 4 shows that the overlap between our method and FLOSS (hypergeometric test, p-value ≈ 0), ORFScore (hypergeometric test, p-value 4*e*-320) and PhyloP (hypergeometric test, p-value 1*e*-32) significant. This shows that our method successfully identifies misannotated lncRNAs by learning relevant features from the lncRNA nucleotide sequences.

**Fig. 4:**
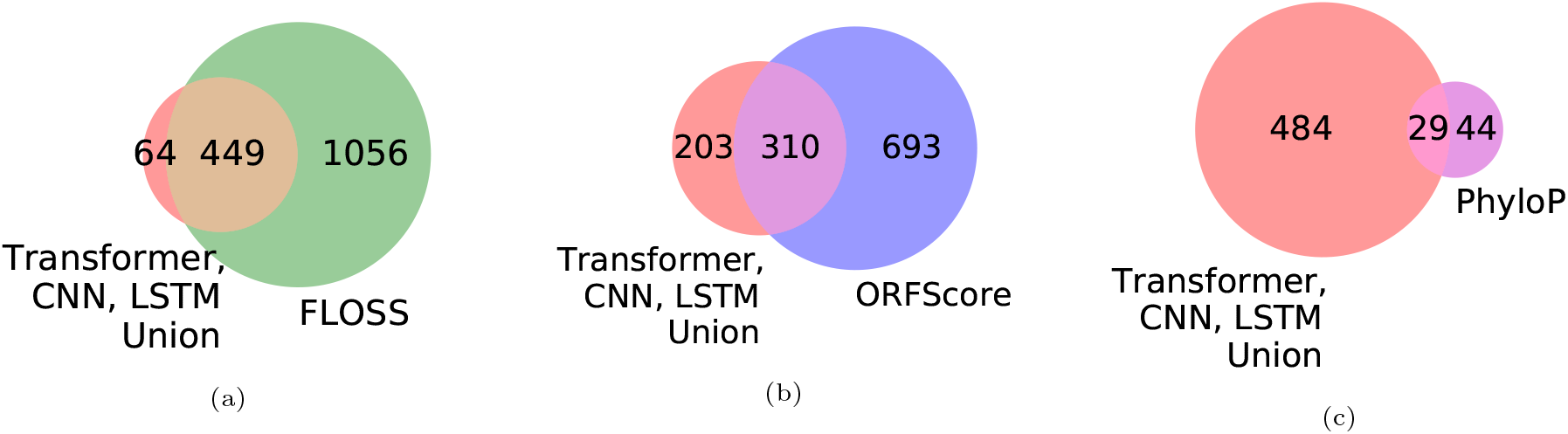
Comparison of misannotated lncRNAs found by CNN, LSTM and Transformer models’ training dynamics with previous ribo-seq data based methods used to find misannotated lncRNAs from: **(a)** FLOSS (p-value *≈* 0), **(b)** ORFScore (p-value 4*e−* 320) and **(c)** PhyloP (p-value (1*e −* 32)) for the dataset from [12]. Background set has 26857 lncRNAs.

### 3.5 Misannotated lncRNAs exist in a continuous cluster with coding RNAs

To analyze the coding and noncoding transcript distributions of the data, we calculated features on for all RNAs in the dataset, based on properties of the transcripts as in [38]. These features include ORF length, ORF quality, nucleotide distribution, translated peptide stability etc. used by [38] (see Table S1 for more details). Using these features, we apply T-distributed stochastic neighbor embedding (t-SNE) [27] (SciKit-learn implementation [34], perplexity=150, iterations=1000, learning rate=200) to reveal RNA clusters.

Figure 5 shows the clusters obtained by performing t-SNE [27] on these features generated from RNA sequences. The labels of the RNAs (coding, non-coding) are not used while generating the clusters. However, based on available coding and non-coding ground-truth labels, along with the biotype information for the ncRNAs, we label each individual RNA example. LncRNAs determined as misannotations by the different deep learning models are labeled in black; interestingly, putative misannotated lncRNAs lie in a cluster contiguous with coding RNAs. This suggests that there is indeed some continuity between coding and lncRNAs in this embedded space and that the categories might not be as mutually exclusive as we believe, which is consistent with recent research discovering that some lncRNAs encode micropeptodes [13]. In support of this, there are clusters of non-coding RNAs (labelled Misc RNA) that are well separated from coding RNAs and that do not contain many putative misannotated lncRNAs.

**Fig. 5:**
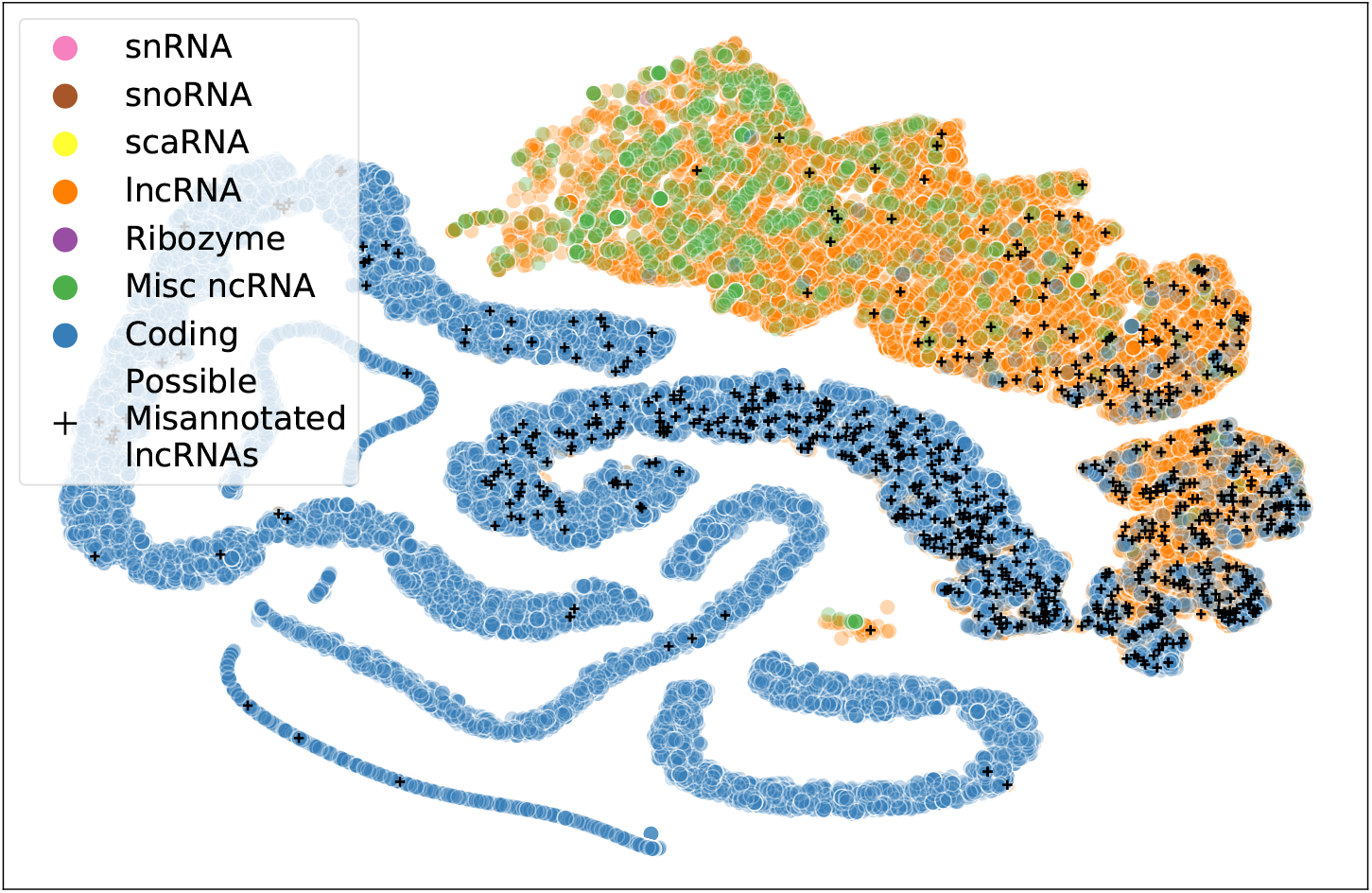
Misannotated lncRNAs exist in a continuous cluster with coding RNAs. t-SNE clusters obtained from hand-crafted features [38] generated from nucleotide sequences. Labels (Coding, lncRNA etc.) are only used for visualizing the clusters, not for generating the clusters. Putative misannotated lncRNAs lie in a cluster contiguous with coding RNAs. There are other clusters of ncRNAs that are well separated from coding RNAs and that do not contain any putative misannotated lncRNAs.

### 3.6 Exploring features learnt by models

To understand which which regions of the sequence are useful for making classification decision, we visualize the activation weights of the model layers. These activation weights determine which sequence features are paid most attention to by the model. Figure 6 shows an example attention map of a misannotated lncRNA generated from the first convolutional layer of the CNN model. Supplementary Figure 1 shows the attention weights visualized for a coding and long non-coding RNA that are not misannotated according to the criteria described above. The CNN model appears to focus on continuous stretches of adenines in the sequence to make decisions about whether a given RNA is coding or long non-coding. This might be because the poly-adenylation sites are one of the major distinguishing features between coding and non-coding RNAs. Supplementary Figure S1 shows the average attention given to all codons for this sequence. Codons with high ‘adenine’ content have higher average attention, but codons ending with ‘TA’ like ‘ATA’, ‘CTA’, ‘GTA’ & ‘TTA’ also have high average attention. Studying these and comparing the average attention differences in codons between coding and non-coding RNAs might prove interesting.

**Fig. 6:**
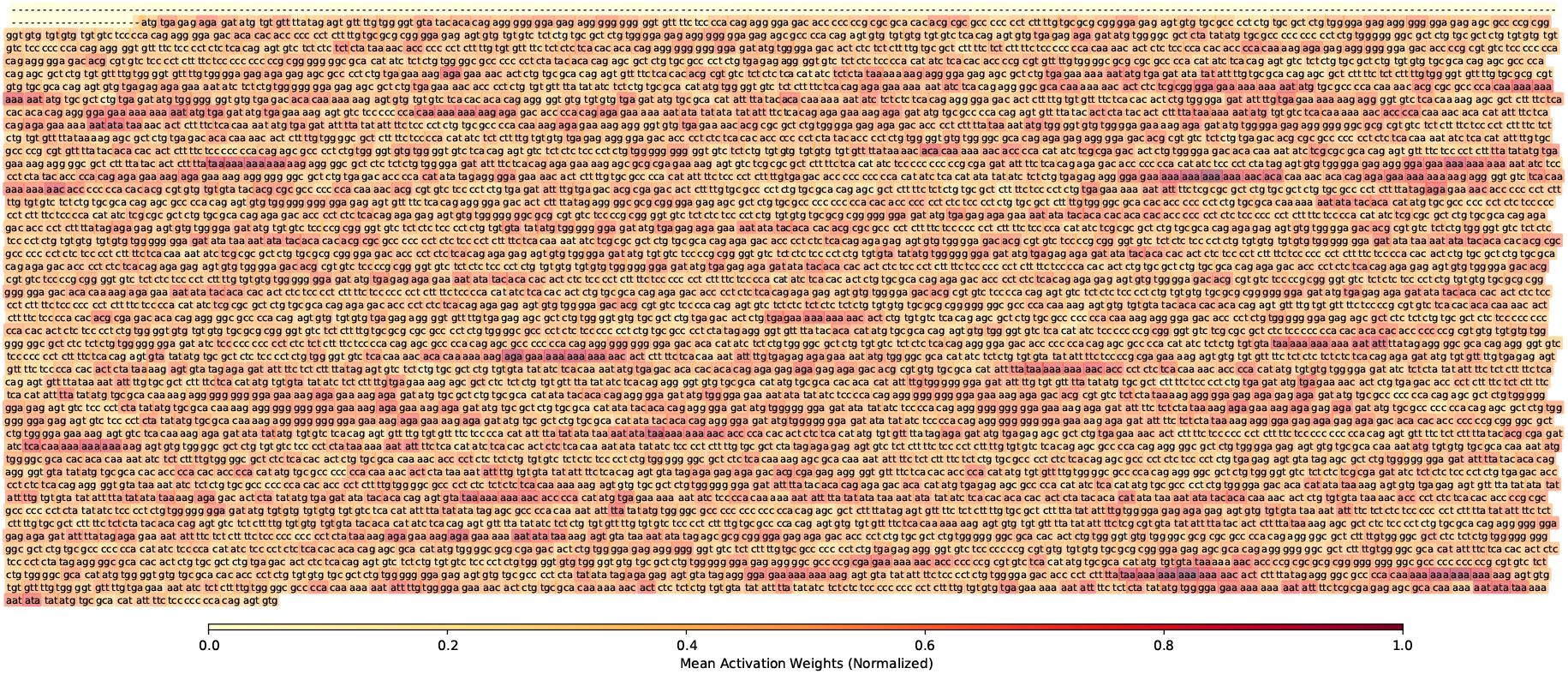
Attention maps explain which parts of sequence are important for making a classification decision. An example attention map extracted from first convolutional layer of CNN model for ENST00000447563 (a misannotated lncRNA). The ground-truth for this RNA is ‘Non-coding’. However, CNN model predicts that this is a non-coding RNA with probability 0.12. Attention visualization shows regions with contiguous ‘A’ nucleotides have high activation weights.

## 4 Conclusion and Future Work

In this paper, we apply the general framework described by [36] for detecting mislabelled samples in a training dataset to detect misannotated lncRNAs. The training dataset, comprising of nucleotide sequences of coding and non-coding RNA, is used to train CNN, LSTM and Transformer models. At the end of each training epoch, coding and non-coding prediction probabilities for every RNA sample are saved. Calculating the mean and standard deviation of the ground-truth class helps determine whether a given RNA is possibly mislabelled. LncRNAs with low mean and standard deviation for the non-coding class are the possible misannotations.

A large number of misannotated lncRNAs are identified by all three different deep learning methods. This is significant since the algorithms to distinguish between coding and non-coding RNAs employed by the models are different. Moreover, when we compare the misannotations discovered here to previous methods to detect misannotated lncRNAs from ribo-seq data and manually curated dataset, we see a large overlap for two of the methods, suggesting that our method is successfully able to detect misannotated lncRNAs. It is also interesting to note that our method shows a high overlap with a manually curated list of misannotated lncRNAs. Therefore, we conclude that this approach offers promising potential for use in curating datasets used for training coding potential predictors and assisting experimental efforts in characterization of misannotated lncRNAs.

To our knowledge, this work also represents the first instance in which nucleotide sequence embeddings and transformer models are applied to the problem of building coding potential predictive models. Using nucleotide embeddings might be preferable to other representations like one-hot encoding [14] or integer encoding [6] used previously. This is because embeddings are learnt from the complete human genome and incorporate the context in which a given codon is found in the DNA. All models are configured to have trainable embeddings; this helps to learn better representations of codons in RNA, since the original embeddings are learnt from DNA sequences. Future work to compare the original embeddings to the embeddings generated from models trained here might provide valuable insight into the differences between codons in DNA and RNA.

One limitation of the approach presented here is that it is computationally intensive since models need to be retrained on the complete dataset after evaluation of the test set performance. However, this approach represents the first method that can find possibly misannotated lncRNAs from the nucleotide sequence alone. In conjugation with ribo-seq data, it can be used to identify misannotated lncRNAs with high confidence. Moreover, it can be used for curating the training datasets used for training coding potential predictors. Future work that compares the misannotated lncRNAs obtained from models here with ribo-seq datasets from different cell-types will provide interesting results on the cell-line specificity of misannotated lncRNAs.

## References

1. Andrews, S.J., Rothnagel, J.A.: Emerging evidence for functional peptides encoded by short open reading frames. Nature Reviews Genetics 15(3), 193–204 (2014)

2. Baek, J., Lee, B., Kwon, S., Yoon, S.: Lncrnanet: long non-coding rna identification using deep learning. Bioinformatics 34(22), 3889–3897 (2018)

3. Batista, P.J., Chang, H.Y.: Long noncoding rnas: cellular address codes in development and disease. Cell 152(6), 1298–1307 (2013)

4. Bazzini, A.A., Johnstone, T.G., Christiano, R., Mackowiak, S.D., Obermayer, B., Fleming, E.S., Vejnar, C.E., Lee, M.T., Rajewsky, N., Walther, T.C., et al.: Identification of small orfs in vertebrates using ribosome footprinting and evolutionary conservation. The EMBO journal 33(9), 981–993 (2014)

5. Brar, G.A., Weissman, J.S.: Ribosome profiling reveals the what, when, where and how of protein synthesis. Nature reviews Molecular cell biology 16(11), 651–664 (2015)

6. Camargo, A.P., Sourkov, V., Pereira, G.A.G., Carazzolle, M.F.: Rnasamba: neural network-based assessment of the protein-coding potential of rna sequences. NAR Genomics and Bioinformatics 2(1), lqz024 (2020)

7. Choi, S.W., Kim, H.W., Nam, J.W.: The small peptide world in long noncoding rnas. Briefings in bioinformatics 20(5), 1853–1864 (2019)

8. Chollet, F., et al.: Keras. blue https://keras.io (2015)

9. Couso, J.P., Patraquim, P.: Classification and function of small open reading frames. Nature reviews Molecular cell biology 18(9), 575 (2017)

10. Derrien, T., Johnson, R., Bussotti, G., Tanzer, A., Djebali, S., Tilgner, H., Guernec, G., Martin, D., Merkel, A., Knowles, D.G., et al.: The gencode v7 catalog of human long noncoding rnas: analysis of their gene structure, evolution, and expression. Genome research 22(9), 1775–1789 (2012)

11. Djebali, S., Davis, C.A., Merkel, A., Dobin, A., Lassmann, T., Mortazavi, A., Tanzer, A., Lagarde, J., Lin, W., Schlesinger, F., et al.: Landscape of transcription in human cells. Nature 489(7414), 101–108 (2012)

12. Elkon, R., Loayza-Puch, F., Korkmaz, G., Lopes, R., Van Breugel, P.C., Bleijerveld, O.B., Altelaar, A.M., Wolf, E., Lorenzin, F., Eilers, M., et al.: Myc coordinates transcription and translation to enhance transformation and suppress invasiveness. EMBO reports 16(12), 1723–1736 (2015)

13. Hartford, C.C.R., Lal, A.: When long noncoding becomes protein coding. Molecular and Cellular Biology 40(6) (2020)

14. Hill, S.T., Kuintzle, R., Teegarden, A., Merrill III, E., Danaee, P., Hendrix, D.A.: A deep recurrent neural network discovers complex biological rules to decipher rna protein-coding potential. Nucleic acids research 46(16), 8105–8113 (2018)

15. Hochreiter, S., Schmidhuber, J.: Long short-term memory. Neural computation 9(8), 1735–1780 (1997)

16. Huang, Y., Wang, J., Zhao, Y., Wang, H., Liu, T., Li, Y., Cui, T., Li, W., Feng, Y., Luo, J., et al.: cncrnadb: a manually curated resource of experimentally supported rnas with both protein-coding and noncoding function. Nucleic Acids Research (2020)

17. Ingolia, N.T.: Ribosome profiling: new views of translation, from single codons to genome scale. Nature Reviews Genetics 15(3), 205–213 (2014)

18. Ingolia, N.T., Ghaemmaghami, S., Newman, J.R., Weissman, J.S.: Genome-wide analysis in vivo of translation with nucleotide resolution using ribosome profiling. science 324(5924), 218–223 (2009)

19. Ingolia, N.T., Lareau, L.F., Weissman, J.S.: Ribosome profiling of mouse embryonic stem cells reveals the complexity and dynamics of mammalian proteomes. Cell 147(4), 789–802 (2011)

20. Ji, Z., Song, R., Regev, A., Struhl, K.: Many lncrnas, 5’utrs, and pseudogenes are translated and some are likely to express functional proteins. elife 4, e08890 (2015)

21. Kang, Y.J., Yang, D.C., Kong, L., Hou, M., Meng, Y.Q., Wei, L., Gao, G.: Cpc2: a fast and accurate coding potential calculator based on sequence intrinsic features. Nucleic acids research 45(W1), W12–W16 (2017)

22. Kingma, D.P., Ba, J.: Adam: A method for stochastic optimization. arXiv preprint arXiv:1412.6980 (2014)

23. Kong, L., Zhang, Y., Ye, Z.Q., Liu, X.Q., Zhao, S.Q., Wei, L., Gao, G.: Cpc: assess the protein-coding potential of transcripts using sequence features and support vector machine. Nucleic acids research 35(suppl 2), W345–W349 (2007)

24. LeCun, Y., Boser, B., Denker, J.S., Henderson, D., Howard, R.E., Hubbard, W., Jackel, L.D.: Backpropagation applied to handwritten zip code recognition. Neural computation 1(4), 541–551 (1989)

25. Li, L., Jamieson, K., DeSalvo, G., Rostamizadeh, A., Talwalkar, A.: Hyperband: A novel bandit-based approach to hyperparameter optimization. The Journal of Machine Learning Research 18(1), 6765–6816 (2017)

26. Lu, S., Zhang, J., Lian, X., Sun, L., Meng, K., Chen, Y., Sun, Z., Yin, X., Li, Y., Zhao, J., et al.: A hidden human proteome encoded by ‘non-coding’genes. Nucleic acids research 47(15), 8111–8125 (2019)

27. Maaten, L.v.d., Hinton, G.: Visualizing data using t-sne. Journal of machine learning research 9(Nov), 2579–2605 (2008)

28. Matsumoto, A., Pasut, A., Matsumoto, M., Yamashita, R., Fung, J., Monteleone, E., Saghatelian, A., Nakayama, K.I., Clohessy, J.G., Pandolfi, P.P.: mtorc1 and muscle regeneration are regulated by the linc00961-encoded spar polypeptide. Nature 541(7636), 228–232 (2017)

29. Miller, W., Rosenbloom, K., Hardison, R.C., Hou, M., Taylor, J., Raney, B., Burhans, R., King, D.C., Baertsch, R., Blankenberg, D., et al.: 28-way vertebrate alignment and conservation track in the ucsc genome browser. Genome research 17(12), 1797–1808 (2007)

30. Nelson, B.R., Makarewich, C.A., Anderson, D.M., Winders, B.R., Troupes, C.D., Wu, F., Reese, A.L., McAnally, J.R., Chen, X., Kavalali, E.T., et al.: A peptide encoded by a transcript annotated as long noncoding rna enhances serca activity in muscle. Science 351(6270), 271–275 (2016)

31. Ng, P.: dna2vec: Consistent vector representations of variable-length k-mers. arXiv preprint arXiv:1701.06279 (2017)

32. Olexiouk, V., Van Criekinge, W., Menschaert, G.: An update on sorfs. org: a repository of small orfs identified by ribosome profiling. Nucleic acids research 46(D1), D497–D502 (2018)

33. O’Malley, T., Bursztein, E., Long, J., Chollet, F., Jin, H., Invernizzi, L., et al.: Keras Tuner. blue https://github.com/keras-team/keras-tuner (2019)

34. Pedregosa, F., Varoquaux, G., Gramfort, A., Michel, V., Thirion, B., Grisel, O., Blondel, M., Prettenhofer, P., Weiss, R., Dubourg, V., Vanderplas, J., Passos, A., Cournapeau, D., Brucher, M., Perrot, M., Duchesnay, E.: Scikit-learn: Machine learning in Python. Journal of Machine Learning Research 12, 2825–2830 (2011)

35. Rinn, J.L., Chang, H.Y.: Genome regulation by long noncoding rnas. Annual review of biochemistry 81, 145–166 (2012)

36. Swayamdipta, S., Schwartz, R., Lourie, N., Wang, Y., Hajishirzi, H., Smith, N.A., Choi, Y.: Dataset cartography: Mapping and diagnosing datasets with training dynamics (2020)

37. Tong, X., Hong, X., Xie, J., Liu, S.: Cppred-sorf: Coding potential prediction of sorf based on non-aug. BioRxiv (2020)

38. Tong, X., Liu, S.: Cppred: coding potential prediction based on the global description of rna sequence. Nucleic acids research 47(8), e43–e43 (2019)

39. Ulitsky, I., Bartel, D.P.: lincrnas: genomics, evolution, and mechanisms. Cell 154(1), 26–46 (2013)

40. Vaswani, A., Shazeer, N., Parmar, N., Uszkoreit, J., Jones, L., Gomez, A.N., Kaiser, L., Polosukhin, I.: Attention is all you need. In: Advances in neural information processing systems. pp. 5998–6008 (2017)

41. Wang, L., Park, H.J., Dasari, S., Wang, S., Kocher, J.P., Li, W.: Cpat: Coding-potential assessment tool using an alignment-free logistic regression model. Nucleic acids research 41(6), e74–e74 (2013)

42. Zhu, M., Gribskov, M.: Mipepid: Micropeptide identification tool using machine learning. BMC bioinformatics 20(1), 559 (2019)

