## Supplementary for "Detecting Misannotated Long Non-coding RNAs with Training Dynamics of Deep Sequence Classification"

#### 1 Features generated from all ncRNAs to use as input for t-SNE clustering

Table S1: List of hand-crafted features used to train machine learning model

| Property | Description | Number of features |
|---|---|---|
| ORF length | length of the longest possible ORF | 1 |
| ORF coverage | quality of ORF | 1 |
| Fickett score | codon bias for 4 nucleotides | 1 |
| Hexamer score | hexamer usage bias | 1 |
| ORF integrity | Binary, whether ORF contains start and stop codon | 1 |
| Isoelectric point | pH at which molecule carries no net charge | 1 |
| Gravy | average hydropathicity of predicted peptide | 1 |
| Instability | estimated stability of predicted peptide | 1 |
| Composition | percentage of each of the 4 nucleotides | 4 |
| Transition | percent frequency of transition from each of nt to other nts | 6 |
| Distribution | distribution for each nt 25% intervals along sequence | 20 |

#### 2 Attention visualization

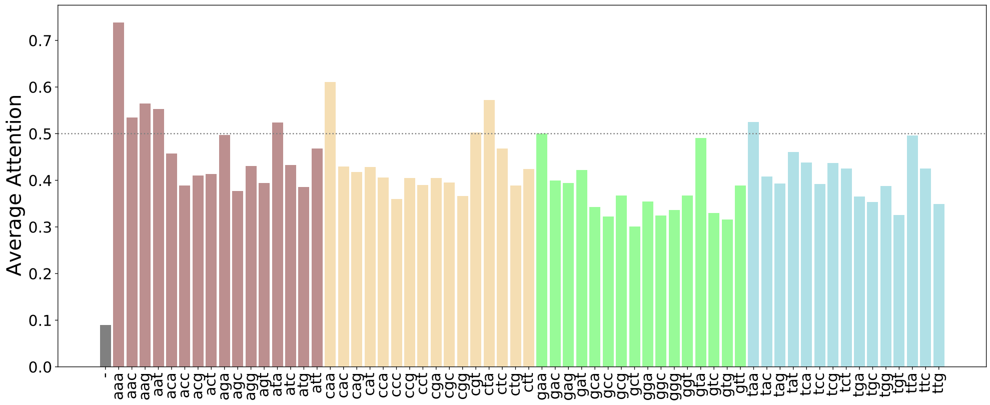


(a)

Figure S1: Attention per codon for ENST00000447563, generated from CNN model.

### 3 Methods to determine true positives from Ribo-seq data: FLOSS, ORFScore and PhyloP

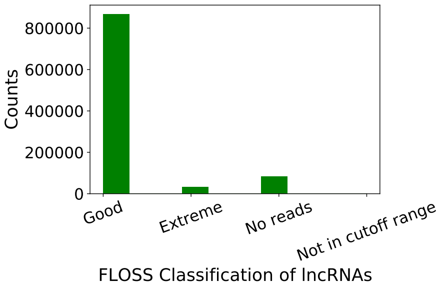


(a)

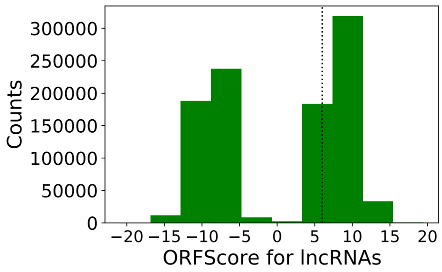


(b)

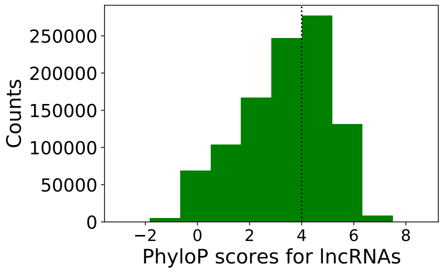


(c)

Figure S2: FLOSS, ORFScore and PhyloP distributions for lncRNAs for ribo-seq data from the U2OS cell-line generated by [1]. The scores are downloaded from sORFs.org [2].

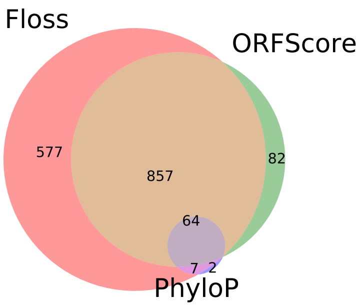


(a)

Figure S3: Overlap between the putative misannotated lncRNAS discovered by FLOSS, ORF-Score and PhyloP for ribo-seq data generated from U2OS cell-line by [1]

### References

[1] Elkon, R., Loayza-Puch, F., Korkmaz, G., Lopes, R., Van Breugel, P.C., Bleijerveld, O.B., Altelaar, A.M., Wolf, E., Lorenzin, F., Eilers, M., et al.: Myc coordinates transcription and translation to enhance transformation and suppress invasiveness. EMBO reports **16**(12), 1723–1736 (2015)

[2] Olexiouk, V., Van Criekinge, W., Menschaert, G.: An update on sorfs. org: a repository of small orfs identified by ribosome profiling. Nucleic acids research **46**(D1), D497–D502 (2018)
